# Reinforced polymer–nanoparticle hydrogels for subcutaneous and sustained delivery of trastuzumab

**DOI:** 10.1101/2024.07.30.605810

**Authors:** Giovanni Bovone, Stéphane Bernhard, Guillaume Jacquot, Vincent Mittelheisser, Céline Mirjolet, Elia A. Guzzi, Lorenza Garau Paganella, Luca Liebi, Olivier Lefebvre, Jacky Goetz, Loïc Charbonnière, Sébastien Harlepp, Xavier Pivot, Mark W. Tibbitt, Alexandre Detappe

## Abstract

In oncology, the advent of monoclonal antibody (mAbs) therapeutics represents a major breakthrough in various cancer diseases. However, these biotherapies often necessitate iterative hospital visits for intravenous infusion that can alter patient’s quality of life and contribute to the chronic saturation of hospitals. Interestingly, subcutaneous formulations of various mAbs offer a promising alternative facilitating faster administration compared with traditional intravenous methods, while still maintaining the same dosing schedule and providing time-saving advantages. Here, we developed an injectable mAb delivery platform using α-cyclodextrin (αCD)-reinforced polymer–nanoparticle hydrogels to perform a subcutaneous injection but also to delay the release of mAbs. By leveraging the versatility of our platform, we formulated hyaluronic acid- and alginate-based injectable drug depots by simply mixing components that are generally regarded as safe (GRAS). We used trastuzumab for the polymer–antibody complexation. The hydrogel depots delayed mAb release up to at least 3 days in both in vitro and in vivo mice models, outperforming clinically approved Herceptin subcutaneous formulation composed of trastuzumab with recombinant human hyaluronidase (rHuPH20).

## Introduction

Monoclonal antibodies (mAbs) have emerged as a groundbreaking advancement in the field of oncology, revolutionizing cancer treatments. These mAbs represent a paradigm shift in precision medicine, offering a targeted and tailored approach to cancer therapy that have improved patient prognosis and survival rates for various cancers. Biotherapies, such as trastuzumab for HER2-positive breast cancer, is a prime example of how mAbs have transformed the treatment landscape. Typically, trastuzumab administration requires regular out-patient visits, either every week or every three weeks, to maintain the therapeutic concentration of the pharmacological active agent over one year or more, depending on the disease stage. These iterative injections have a detrimental impact on patient quality of life. In this context, subcutaneous (SC) formulation of trastuzumab presents a notable advancement, offering significant time-saving benefits in the clinic compared with traditional intravenous (IV) administration. Exploring extended intervals between treatments could further alleviate the negative impact of frequent injections, ultimately improving patient well-being.

The SC route, while beneficial, introduces significant complexities. The architecture of the skin comprises three distinct layers: the epidermis, dermis, and hypodermis. The hypodermis, also referred to as SC tissue, is a loosely organized connective tissue filled with adipocytes, fibroblasts, and immune cells within a vascularized collagenous extracellular matrix (ECM). This ECM forms a structured lattice of collagen and glycosaminoglycans, such as hyaluronic acid and chondroitin sulfate, suspended in interstitial fluid. Hyaluronic acid, with its negative charge, significantly influences the transport dynamics and stability of drugs and excipients within the SC space, particularly affecting biotherapeutic agents like proteins and mAbs. Consequently, the dense networks of blood and lymphatic capillaries, along with the low proteolytic activity within SC tissue, make SC administration an attractive route for pharmaceutical and drug delivery systems. However, compared to IV administration, SC injections result in a slower onset of action as the drug must diffuse from the SC tissue into the bloodstream. Additionally, the SC route is limited by injection volume, as volumes exceeding 2–3 mL can cause pain, discomfort, site leakage, or tissue distortion due to the dense fibrous nature of SC tissue. The restricted fluid flow within the ECM, caused by the thickness of the SC tissue and the dense, negatively charged hyaluronic acid network, impairs drug transport to systemic circulation, leading to slow absorption rates (typically a few milliliters per hour) and incomplete bioavailability, which ranges from 9% to 90%, with a typical trend between 50% and 80%. Furthermore, the unique architecture and composition of the SC space significantly influence the fate of nanoscale drug delivery systems administered via the SC route, contributing to heterogeneity in patient when compared to the direct IV route.

To address the challenge posed by the ECM barrier and enable the administration of larger volumes, various strategies utilizing enzymatic combinations have been explored. Among these, only one has achieved clinical approval, the recombinant human hyaluronidase (rHuPH20) from Halozyme, registered in 2005. This enzyme facilitates SC delivery by depolymerizing hyaluronic acid within the ECM, creating local space for up to 20 mL and thereby enhancing the absorption of the injected product. Nevertheless, these innovations have not provided controlled or sustained release of the mAbs for long-lasting delivery. Hence, to pave this route, several injectable biomaterials have been developed to form long-lasting and/or local depots for the sustained release of mAbs, and in particular trastuzumab.^1–5^ Vitamin E-functionalized block copolymers have been used to create micellar hydrogel networks that facilitate the release of trastuzumab in a murine model.^1,2^ Hyaluronic acid (HA)-based scaffolds exhibited a delayed trastuzumab release, likely due to the formation of polyanion–antibody complexes between the negatively-charged HA and the positively-charged trastuzumab.^3–5^ For many of the systems mentioned, complicated processing steps, such as enzymatic or photo-cross-linking chemistries and microfluidic handling, were required for hydrogel formation. Generally, materials assembled from unmodified components that are approved by the US Federal Drug Administration (FDA) or generally regarded as safe (GRAS) have improved chances of being translated clinically.

Polymer–nanoparticle (PNP) hydrogels comprise a class of injectable biomaterials that form via intermolecular interactions between GRAS-listed polymers and biodegradable nanoparticles (NPs).^6^ PNP hydrogels are injectable through narrow gauge syringes and catheters as they can flow upon applied stress (shear-thin) and reform a stable network (self-heal) when the stresses are removed.^7,8^ They have been applied as drug delivery systems for several applications, including immune cell recruitment, to release COVID-19 vaccines, to deliver immunostimulatory antibodies, and to supply CAR-T cells.^8–11^ However, one limitation in the design of this class of materials for injectable drug delivery applications is their limited range of mechanical properties and restricted design space. Recently, the addition α-cyclodextrin (αCD) to PNP formulations improved the mechanical properties and enabled the utilization of diverse building blocks (both polymers and NPs) for hydrogel formation.^12^

In this work, we leveraged the versatility of αCD-reinforced PNP (CD–PNP) hydrogels to design two injectable formulations: one based on hyaluronic acid (HA) and the other on alginate. We validated the in vitro release profiles of trastuzumab for both formulations and assessed the stability of trastuzumab post-release from the hydrogel to ensure its integrity remains intact. Furthermore, we performed pharmacokinetic studies in mice where both formulations were administered subcutaneously and compared to both the IV and SC clinically approved formulation of trastuzumab.

## Results and Discussion

### Design of CD–PNP hydrogels for trastuzumab delivery

Trastuzumab is a first-in-class monoclonal antibody for the treatment of HER-positive breast cancer.^13^ However, multiple subcutaneous or intravenous injections are needed to maintain therapeutic antibody levels in the body. This results in frequent hospital visits, which impacts life quality. A sustained release system for trastuzumab (and potentially other therapeutic antibodies) is needed to improve clinical application of these powerful therapeutics. To address this need, we prepared trastuzumab-loaded CD–PNP hydrogels were prepared by mixing PEGylated nanoparticles, αCD, a biopolymer, and trastuzumab (**Figure 1**).^12^ Given the target of in vivo application, we assembled trastuzumab-loaded CD–PNP hydrogels using GRAS building blocks: PEG-*block*-poly(*D,L*-lactide) (PEG-*b*-PLA) NPs, αCD, and pharma-grade alginate or HA (**Figure 1a**). At the molecular scale, the threading of αCD onto the PEG chains of nanoparticles (NPs) and the subsequent formation of polypseudorotaxanes would enhance the interactions between NPs, improving the mechanical properties of the injectable biomaterials.^14^ To ensure biocompatibility, we used HA as the structural polymer given its natural presence in connective, epithelial, and neural tissues. Since HA cannot undergo cross-linking without chemical modification, we selected alginate as an alternative structural polymer to enable an optional post-stabilization by cross-linking with divalent cations (Ca^2+^).

**Figure 1.**
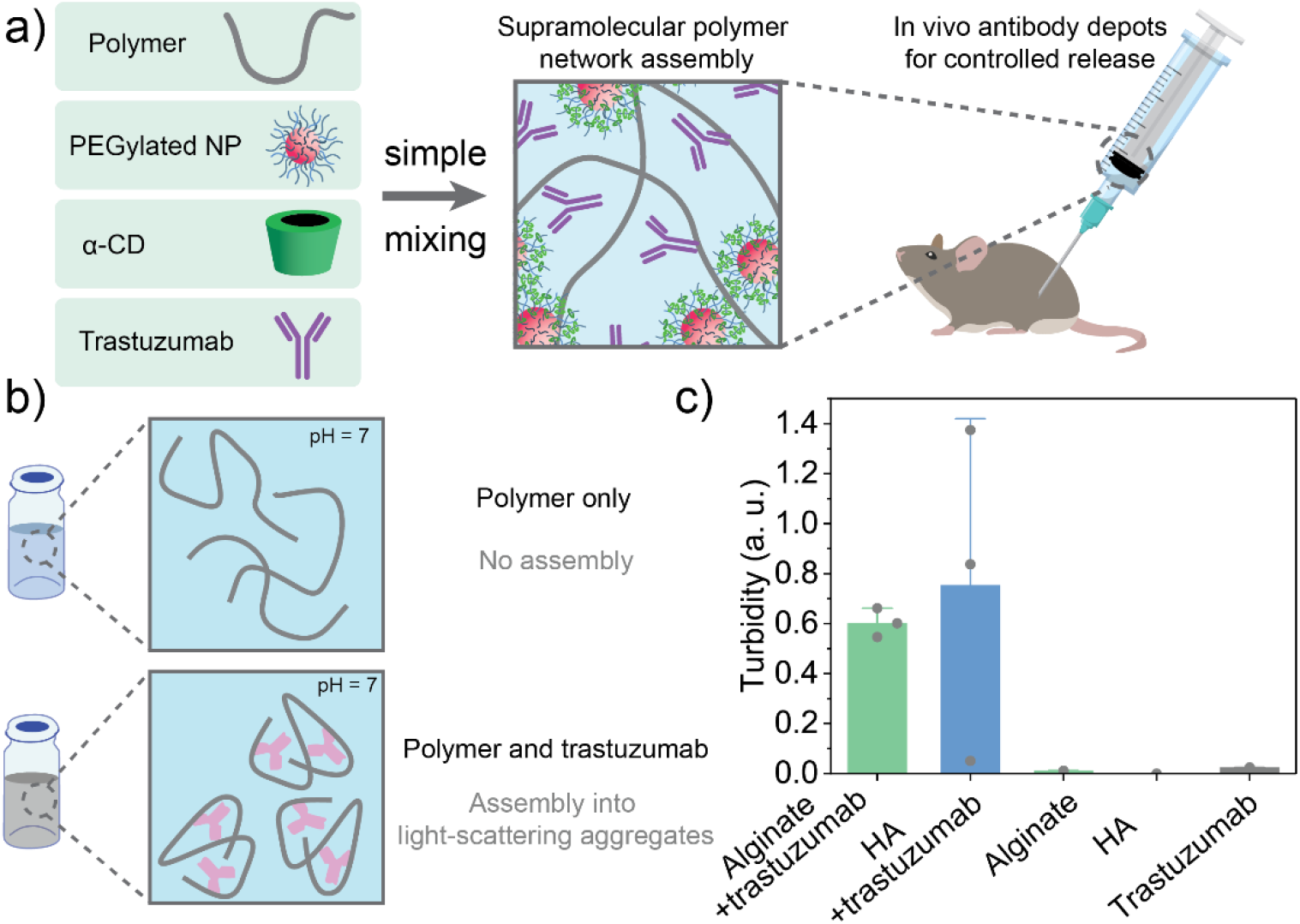
CD–PNP hydrogels for the controlled release of trastuzumab in vivo. a) Trastuzumab-encapsulating CD–PNP hydrogels were formed by simply mixing αCD, PEGylated NPs, alginate or hyaluronic acid (HA), and trastuzumab. Network formation was promoted by threading of αCD onto NP PEG chains and subsequent polypseudorotaxane formation. The designed formulations were used for in vivo controlled release of trastuzumab. b) At physiological pH, alginate and HA are negatively-charged, whereas trastuzumab is positively-charged.^3^ This is advantageous as it allows complexation of the trastuzumab with the support polymers during gel preparation, providing sustained release.^5^ c) Turbidity measured at 450 nm supports that aggregates are formed upon mixing alginate and trastuzumab.

Trastuzumab was encapsulated into CD–PNP hydrogels via incorporation within the hydrogel precursors. Given that the pKa is ∼9.2 for trastuzumab, ∼5.4 for alginate, and ∼2.5 for HA, ^15–17^ trastuzumab is positively-charged at physiological pH (∼7.4) whereas alginate and HA are negatively-charged. This facilitates the formation of polymer–trastuzumab complexes that can prolong the release of mAbs for both formulations (**Figure 1b**): HA CD–PNP (2 wt% HA, 5 wt% NPs, 10 wt% αCD) and Alg CD–PNP (2 wt% alginate, 5 wt% NPs, 10 wt% αCD).^3–5^ At physiological pH, an increase in turbidity was observed when alginate and trastuzumab were mixed (**Figure 1c**). The turbidity of solely alginate or trastuzumab were similar to the one of PBS only. This suggested assembly of trastuzumab with the structural polymers used in our formulations.^5^

### CD–PNP hydrogels were moldable and allowed post-fabrication stabilization

The rheological properties of CD–PNP hydrogels were characterized via shear rheometry (**Figure 2**). The frequency behavior of alginate and HA CD–PNP formulations was described with oscillatory frequency sweeps in the linear viscoelastic region (*γ* = 0.1 %; **Figure S1a**). We first characterized CD–PNP formulations that were assembled solely via non-covalent interactions between the polymers, αCD, and the NPs.

**Figure 2.**
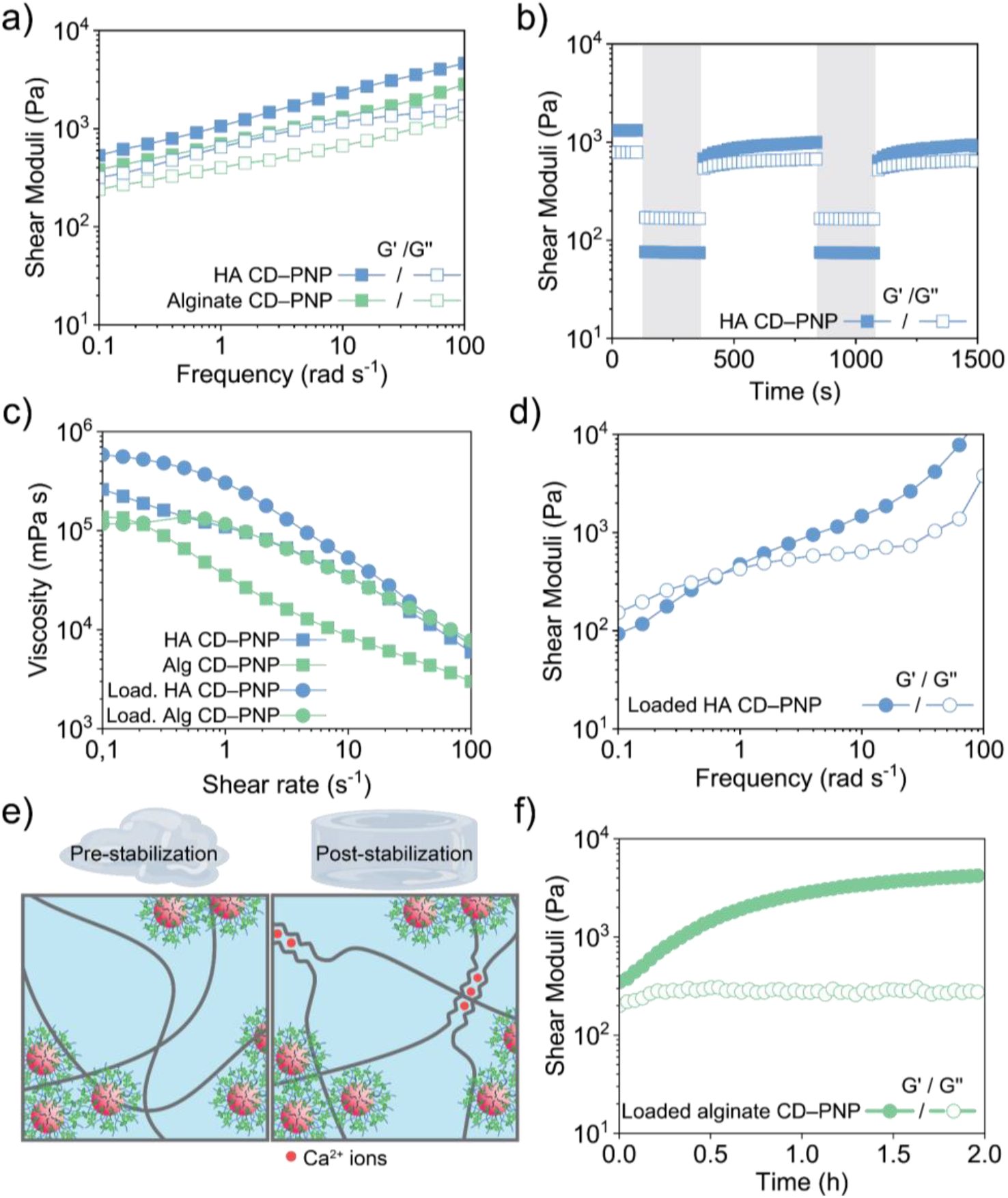
HA and alginate CD–PNP hydrogels are suitable scaffolds to make antibody depots. a) Oscillatory frequency sweep of HA CD–PNP and alginate CD–PNP hydrogels (*γ* = 0.1 %). Both formulations formed solid-like viscoelastic hydrogels over the whole frequency range (*G’* > *G’’*). b) HA CD–PNP hydrogels were subjected to periods of high oscillatory shear strains (*γ* = 200 %; *ω* = 10 rad s^-1^) alternated to periods of low oscillatory shear strains (*γ* = 0.1 %; *ω* = 10 rad s^-1^) to simulate syringe extrusion. At high shear strains, the hydrogel network was broken (*G’* < *G’’*) and partially recovered during low strain periods (*G’* > *G’’*). c) Rotational shear rate ramp experiments of trastuzumab-loaded and non-loaded CD– PNP formulations demonstrate a shear-thinning behavior. d) Oscillatory frequency sweeps of HA CD–PNP loaded with 12 wt% monoclonal antibody trastuzumab (*γ* = 0.1 %). The data suggested an acceleration of polymer dynamics. e) Alginate CD–PNP hydrogels allowed post-fabrication stabilization via ionic Ca^2+^ crosslinking. f) Time sweeps of trastuzumab-loaded alginate CD–PNP showed that Ca^2+^ post-stabilization occurred within ∼2 h (*γ* = 0.1 %; *ω* = 10 rad s^-1^).

Both HA CD–PNP and alginate CD–PNP hydrogels had storage moduli, *G’*, between 0.1 kPa and 1 kPa and solid-like properties (*G’* > *G’’*) over the entire frequency spectrum tested (*ω* = 0.1 – 100 rad s^-1^, **Figure 2a**). The moduli increased with increasing frequency, consistent with the presence of entangled polymers.

The self-healing (elastic recovery) behavior of CD–PNP formulations was studied by alternating high oscillatory shear strain intervals (*γ* = 200 % or 10 %, *ω* = 10 rad s^-1^) and low oscillatory shear strain intervals (*γ* = 0.1 %, *ω* = 10 rad s^-1^; **Figure 2b** and **Figure S1b**). At high shear the hydrogel network was broken (*G’* < *G’’*) and, at low shear, the solid-like properties were partially recovered (*G’* > *G’’*). We characterized how loading clinically-relevant amounts of trastuzumab (120 mg mL^-1^) impacts the hydrogel rheological properties. Rotational shear rate ramp experiments showed that all non-loaded and loaded HA CD–PNP and alginate CD–PNP hydrogels were shear-thinning (**Figure 2c**). We fitted the viscosity versus shear rate data between 1 and 100 s^-1^ to a power law model 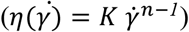 and found that all hydrogels had *n* < 0.5 confirming the shear-thinning behavior. Frequency sweeps of HA CD–PNP hydrogels loaded with trastuzumab showed solid-like behavior for *ω* > 1 rad s^-1^ (**Figure 2d**). The yield strains of non-loaded hydrogels were ∼5 % for alginate CD–PNP and ∼10 % for HA CD–PNP (**Figure S1a**). After trastuzumab loading, the yield strain increased to ∼100 % for HA CD–PNP (**Figure S1c**) and decreased to < 1 % for alginate CD–PNP (**Figure S1d**). Since antibody loading interfered with the stability of the hydrogel network (**Figure 2e, Figure S1e**), we further investigated scaffold post-stabilization strategies by further cross-linking alginate with Ca^2+^ ions (**Figure 2f** and **Figure S1f**). We included solid CaCO_3_ and glucono-*delta*-lactone (GDL; a cyclic ester) into the formulation. The slow hydrolysis of GDL reduces the local pH of the solution. This promotes a gradual dissolution of CaCO_3_ releasing the Ca^2+^ ions necessary for post-crosslinking the network.^18^ In alginate CD–PNP hydrogels, released Ca^2+^ ions progressively cross-linked and stabilized the network (**Figure 2f**). Time-sweeps of trastuzumab-loaded alginate CD–PNP hydrogels indicated that the final storage modulus was reached within hours (*G’* ∼ 1.4 kPa; *γ* = 0.1 %; *ω* = 10 rad s^-1^).

### CD–PNP depots delayed the release of trastuzumab in vitro

We tested the ability of CD–PNP hydrogels to delay the release of trastuzumab by including a clinically-relevant loading of antibody with the hydrogel formulation. In the commercially available subcutaneous formulation, the dosage for humans is ∼600 mg of trastuzumab dissolved into 5 mL of physiological solution (∼12 wt%).^19^ We loaded an equivalent amount of trastuzumab in HA CD–PNP and alginate CD– PNP hydrogels and studied the in vitro drug release properties in phosphate-buffered saline (PBS) (**Figure 3**).

**Figure 3.**
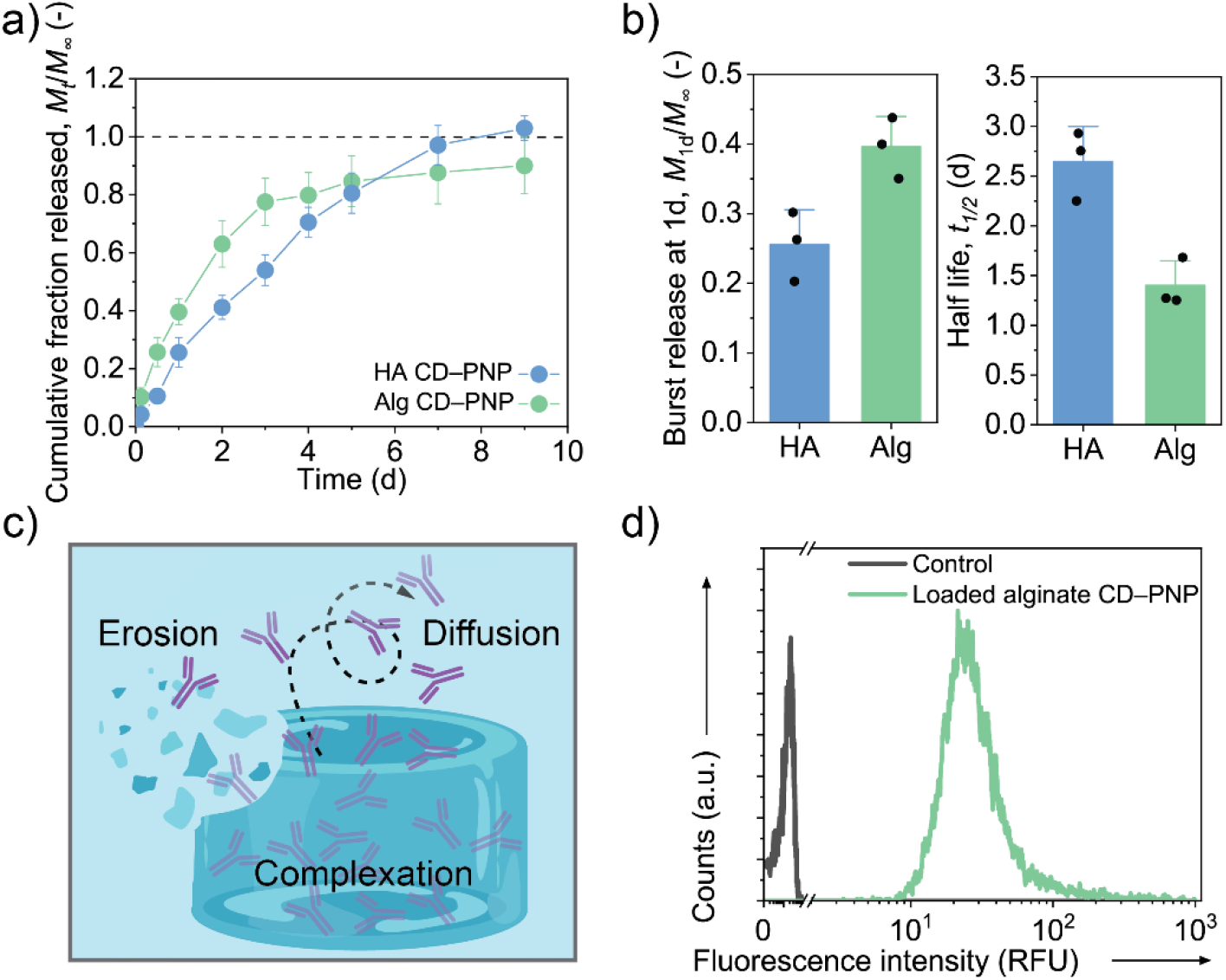
CD–PNP hydrogels delayed the release of trastuzumab in vitro. Trastuzumab (120 mg mL^-1^) was loaded in CD–PNP hydrogels and the amount released in a PBS solution was quantified over time. a) *M*_*t*_/*M*_∞_ represents the cumulative fraction of trastuzumab released up to the timepoint *t* for HA CD–PNP hydrogels (2 wt% HA, 5 wt% NPs, 10 wt% αCD) and alginate CD–PNP hydrogels (3 wt% alginate, 5 wt% NPs, 10 wt% αCD; cross-linked with 0.2 M CaCl_2_). b) Burst release after 24h, *M*_1d_/*M*_∞_, and half-life, *t*_1/2_, of trastuzumab.^20^ Data plotted as mean ± S.D. c) d) Fitting to the Ritger-Peppas model (**Figure S2**) suggested that HA CD–PNP released trastuzumab via an erosion-based mechanism (*n* ∼ 0.85) and alginate CD–PNP via mostly diffusive mechanism (*n* ∼ 0.66).^21,22^ d) Cy5.5-labeled trastuzumab released from CD– PNP hydrogels was incubated with HER2-overexpressing epithelial ductal carcinoma cells (HCC-1954 cells). Flow cytometry analysis demonstrated that the antibody retained the ability to bind to the cells.

We report the data as cumulative fraction of trastuzumab released *M*_*t*_/*M*_*∞*_, where *M*_*t*_ is the mass released up to a specific timepoint, *t*, and *M*_*∞*_ is the total dose loaded (**Figure 3a**). HA CD–PNP hydrogels exhibited an initial burst release of *M*_1d_/*M*_*∞*_ ∼ 0.26, followed by a constant release of trastuzumab up to day 7. Alginate CD–PNP hydrogels had a burst release of *M*_1d_/*M*_*∞*_ ∼ 0.40 and did not fully release their payload over the experimental period of 9 d (**Figure 3b**). The in vitro half-life, *t*_1/2_, is defined as the length of time required for a substance to decrease to half of the initial dose. The half-life of trastuzumab in HA CD–PNP and alginate CD–PNP were *t*_1/2_ ∼ 2.65 d and ∼ 1.40 d respectively. We fitted the first fraction to the Ritger-Peppas model, which describes the Fickian and non-Fickian release behavior of hydrogels,^21,22^ to determine the release exponent, *n*, which is 0.5 for Fickian diffusion, 0.5 < *n* < 1.0 for non-Fickian transport, and *n* ∼ 1 for zero-order release (**Figure S2**). For HA CD–PNP, *n* ∼ 0.85 indicating an erosion-like release mechanism, whereas for alginate CD–PNP, *n* ∼ 0.66 suggesting a diffusion-like mechanism (**Figure 3c**). Trastuzumab diluted in sodium chloride solutions is chemically and physically stable for up to 6 months.^23^

To assess whether hydrogel encapsulation had a detrimental effect on the bioactivity of trastuzumab, we measured the ability of the antibody to bind cellular HER2 and performed cell-based potency assays. We examined binding of fluorescently-labeled trastuzumab on human HER2+ breast cancer cell line (HCC-1954) (**Figure 3d**). Flow cytometry analysis demonstrated an increased fluorescence intensity for cells incubated with the release supernatant from loaded alginate CD–PNP hydrogels in comparison with untreated controls demonstrating that at least a fraction of trastuzumab released from alginate CD–PNP scaffolds retained HER2 binding capability.

Overall, both HA and alginate CD–PNP hydrogels delayed the release of trastuzumab. Since the mesh size of alginate- and HA-based hydrogels is generally much larger than the hydrodynamic diameters of proteins,^3,24,25^ we hypothesized that, initially, trastuzumab was electrostatically complexed with the polymer matrix and was retained within the scaffold. Given the pKa differences of trastuzumab with HA (ΔpKa ∼ 6.7) and alginate (ΔpKa ∼ 3.8), at physiological pH, the data suggested that higher polymer–antibody charge differences corresponded with a decreased burst release (*M*_1d_/*M*_*∞*_ ∼ 0.26 for HA CD–PNP and *M*_1d_/*M*_*∞*_ ∼ 0.40 for alginate CD–PNP).^20^ Fitting to the Ritger-Peppas model suggested that the main elements governing trastuzumab release were an interplay between scaffold erosion and diffusion of non-bound antibodies, where scaffold erosion dominated in HA CD–PNP hydrogels and diffusion of unbound antibodies was more prominent in alginate CD–PNP scaffolds. Trastuzumab that was released from CD– PNP scaffolds maintained the ability to bind human HER2^+^ cancer cells and, given that the scaffolds are composed of GRAS materials, they are suitable antibody depots to be tested in vivo.

### CD–PNP hydrogels effectively and safely localized antibodies in vivo and delayed the release of trastuzumab

Having assessed the delayed antibody release of CD–PNP hydrogels in vitro, we tested if CD–PNP hydrogels were safe to use in vivo (**Figure 4**). As a demonstration of the biocompatibility of CD–PNP hydrogels, initially, CD–PNP hydrogels were injected subcutaneously in female Balb/c mice and compared with control PBS injections. Qualitatively, while the PBS bolus quickly dissipated, the post-stabilized alginate CD–PNP scaffolds remained intact for the entire duration of the experiment (6 d). The injected hydrogels were harvested and the tissues at the injections site were stained for hematoxylin and eosin (**Figure 4a**).

**Figure 4.**
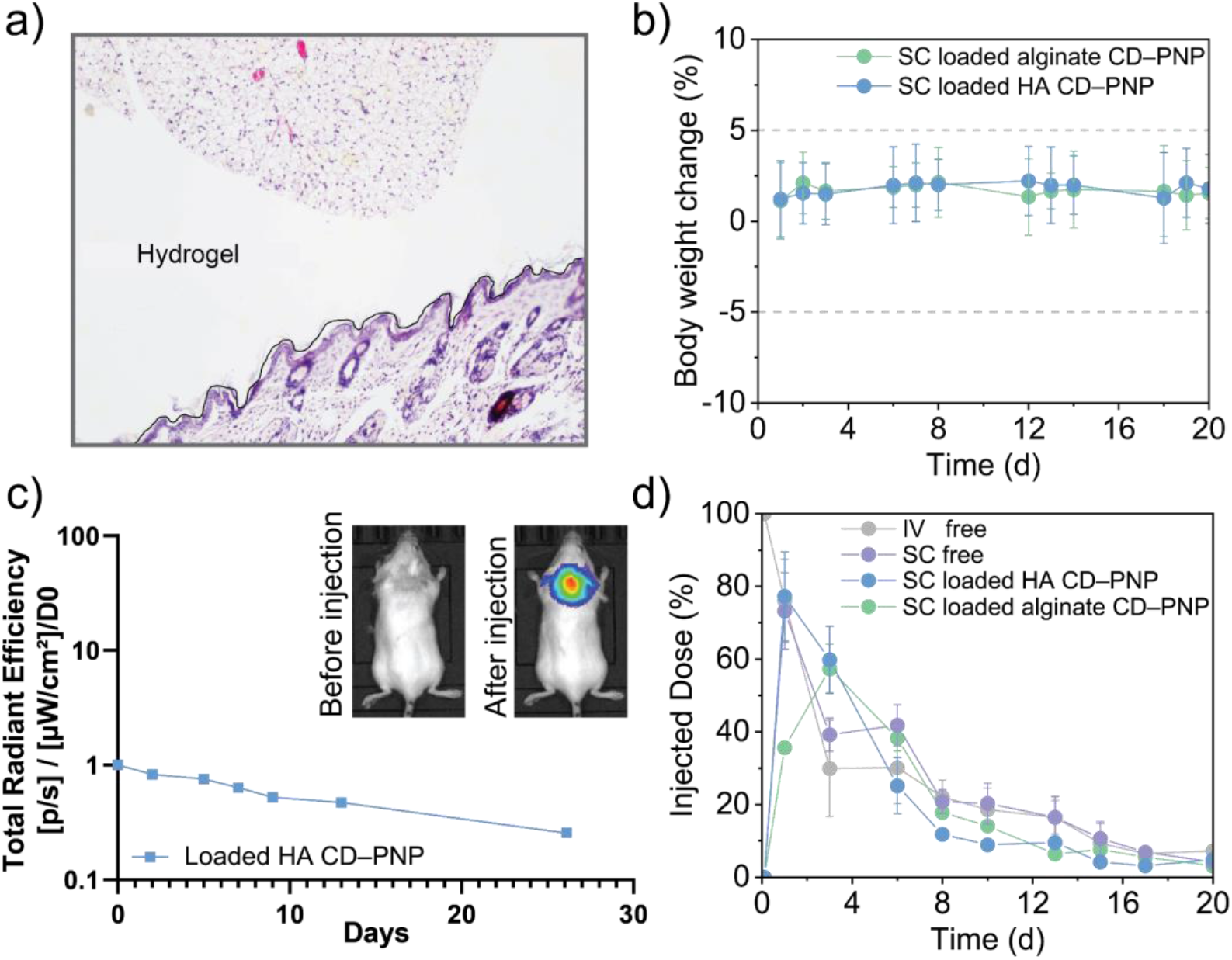
In vivo trastuzumab hydrogel depots for controlled release. **a)** Hematoxylin and eosin staining of mouse injection tissues. The interface between the alginate CD–PNP hydrogel and tissues is marked with a black line. Lymphocytes were present at the interface. No major adverse effects such as fibrosis or damage in adjacent tissues was evident. Cells did not infiltrate into the alginate CD–PNP hydrogel scaffold hinting to reduced scaffold biodegradation. b) The mouse weights remained within ± 5%, indicating no macroscopic adverse effects. Data plotted as mean ± S.E.M. c) HA CD–PNP hydrogels were effective local drug depots of trastuzumab. In vivo fluorescent imaging (n = 1) illustrated the localization of a Cy5.5-labeled trastuzumab in a Balb/c mouse. The antibody was injected subcutaneously with an HA CD–PNP hydrogel depot. Evolution of the total radiant efficiency relatively to day 0. A gradual decrease of radiant efficiency was visible over time. After 26 d, there was still evidence of locally-confined antibody. d) TR-FRET measurements for trastuzumab concentration determination in the blood plasma expressed as a percentage of the injected dose (27.6 mg). Relevant pharmacokinetic parameters are presented in **Table 1**. When compared with free trastuzumab injections, CD–PNP hydrogels delayed the time to reach C_max_ concentration by ∼3 d. Data plotted as mean ± S.D. SC data was retrieved from a dataset containing trastuzumab and rHuPH20.^26^

Lymphocytes concentrated at the interface between the tissue and the scaffold, but no evidence of fibrosis or damage in adjacent muscle tissues was visible over the course of the experiment. No cell infiltration into the scaffolds was visible, suggesting limited remodeling of alginate on this time scale.^27^ We next injected the scaffold subcutaneously and followed the mouse weight for 20 days. The weight of all mice fluctuated between ± 5% (**Figure 4b**) suggesting no macroscopic adverse effects of the injections.

Given that SC CD–PNP injections were not locally immunogenic, we further assessed whether CD–PNP hydrogels could serve as in vivo antibody depots. We subcutaneously injected CD–PNP hydrogels loaded with 120 mg mL^-1^ trastuzumab and 1.25 mg mL^-1^ (∼ 9 µM) of Cy5.5-labeled antibody. In vivo fluorescence imaging corroborated the hypothesis that the antibody remained localized near the site of injection (**Figure 4c**). The total fluorescence decayed over time suggesting a sustained release of trastuzumab.

We next evaluated the pharmacokinetics of trastuzumab following SC administration using the HA CD– PNP and alginate CD–PNP formulations (**Figure 4d**). As controls, we administered the antibody as a liquid solution via IV with PBS as a carrier soltuion and compared with a prior SC dataset containing trastuzumab and rHuPH20.^26^ The relevant pharmacokinetic parameters—maximum concentration in the plasma (*C*_max_), time when *C*_max_ was reached (*t*_max_), half-life (*t*_1/2_), and area under the curve (AUC_0-20d_; **Table 1**) were quantified. The release from CD–PNP hydrogels exhibited a prolonged release profile, similar to the delayed release patterns observed in our in vitro studies.

**Table 1.** Calculated pharmacokinetic parameters for the different formulations. We calculated the primary pharmacokinetic parameters below by converting the relative injected dose to a blood concentration. To do this, we assumed an injected dose of 27.6 mg and 2 mL of total blood volume per mouse. Here, *C*_max_ is maximum blood serum concentration of trastuzumab; *t*_max_ is the time when *C*_max_ was observed; *t*_1/2_ is the half-life; and *AUC*_0-20d_ is the area under the concentration curve.

| Route of administration | Intravenous | Subcutaneous<sup>26</sup> | Subcutaneous | Subcutaneous |
| --- | --- | --- | --- | --- |
| Carrier | PBS | rHuPH20 | HA CD–PNP | Alg CD–PNP |
| $C_{\text{max}}$ | $13.8 \text{ mg mL}^{-1}$ | $10.1 \text{ mg mL}^{-1}$ | $10.6 \text{ mg mL}^{-1}$ | $7.9 \text{ mg mL}^{-1}$ |
| $t_{\text{max}}$ | 0 d | 1 d | 1 d | 3 d |
| $t_{1/2}$ | 2.1 d | 2.4 d | 3.9 d | 4.2 d |
| $\text{AUC}_{0-20\text{d}}$ | $67.7 \text{ mg mL}^{-1} \text{ d}$ | $67.3 \text{ mg mL}^{-1} \text{ d}$ | $57.7 \text{ mg mL}^{-1} \text{ d}$ | $56.8 \text{ mg mL}^{-1} \text{ d}$ |

The *C*_max_ was lower for each SC formulation (*C*_max_ ∼ 7.9–10.1 mg mL^-1^) as compared with the IV injection (*C*_max_ ∼ 14 mg mL^-1^). SC injections delayed *t*_max_ by 1 d^26^ and alginate CD–PNP gels further delayed *t*_max_ to 3 d. While the half-life of trastuzumab in blood sera, *t*_1/2_, for free liquid trastuzumab was between 2.1 and 2.4 d, HA and alginate CD–PNP respectively delayed *t*_1/2_ to 57.7 and 4.2 d demonstrating controlled release. The AUC_0-20d_ of both subcutaneously-injected CD–PNP formulations decreased by 15% when compared to IV and SC injections of liquid trastuzumab solutions. We hypothesize that this difference may be related to antibody retention inside the hydrogels.

Overall, our data demonstrated that the CD–PNP formulations tested in this study were well-tolerated and formed trastuzumab depots for sustained antibody release. When compared with IV injections of liquid trastuzumab formulations, our data confirmed previous literature findings where SC administrations attenuated *C*_max_ and prolonged *t*_1/2_.^19^ Both CD–PNP formulations successfully delayed the release of trastuzumab when compared with IV and SC administration of rHuPH20-trastuzumab. The general decrease of the AUC_0-20d_ with CD–PNP hydrogels might suggest that the antibody was retained in the hydrogel and would have been released at timepoints that go beyond the interval tested in our experiment.^5^ In literature, while many studies related to engineering materials for the controlled release of trastuzumab directly focus on tumor-bearing mice experiments, only few investigate antibody pharmacokinetics.^1–5^ Among them, Kurisawa and coworkers developed an HA–tyramine hydrogel for the sustained release of trastuzumab with *t*_max_ and a *t*_1/2_ comparable to those observed in our research.^3^ Moving forward, we plan to investigate the effect of hyaluronidase on the overall pharmacokinetics since it would (i) partially degrade the extracellular matrix to prevent local trastuzumab entrapment and (ii) break down HA-based hydrogel liberating potentially complexed antibody.

## Conclusion

We designed two locally-injectable CD–PNP hydrogels aimed at delaying the release of antibodies and we studied as lead candidate the sustained release of trastuzumab. At physiological pH, we leveraged electrostatic interactions between positively charged trastuzumab and negatively charged structural polymers, namely HA and alginate, to delay the release of the antibody. Rheological characterization demonstrated that both CD–PNP hydrogel formulations exhibited shear-thinning and self-healing properties. Additionally, the alginate CD–PNP hydrogels allowed post-stabilization of the network by cross-linking with divalent cations. Both HA CD–PNP and alginate CD–PNP delayed the release of trastuzumab by 4 d in vitro. HA CD–PNP hydrogels eroded completely within 6 d releasing the entire load of trastuzumab, whereas alginate CD–PNP hydrogels retained approximately 40% of the loaded trastuzumab inside the network. Preliminary data suggested that hydrogel encapsulation did not impair the capacity of trastuzumab to bind to cellular HER2. Histology and in vivo fluorescence showed that CD– PNP hydrogels might not cause toxicity and could localize trastuzumab in subcutaneous depots. In vivo, when compared with free IV and SC tratuzumab injections, HA CD–PNP and alginate CD–PNP delayed the release of trastuzumab over ∼4 d. Their pharmacokinetic profiles were comparable to SC trastuzumab with rHuPH20.^26^ When compared with IV formulation, the results were comparable in terms of AUC with longer half-life and lower *C*_max_. Overall, our study demonstrated that CD–PNP hydrogels are potential candidates for delivering mAbs in cancer therapies and providing tunable delayed release might allow us to revisit the schedule of iterative administration. Future efforts should be directed at limiting the quantity of residual antibody that remains electrostatically complexed to the hydrogel scaffold and to assess the degradation of the hydrogel.

## Materials and methods

### Materials

Water was deionized, dH_2_O, using a milliQ purification system and a BioPak filter. For in vitro experiments, αCD (Ref. C0776) was purchased from TCI chemicals (Tokyo, Japan). For in vivo experiments, US pharma-grade αCD (Ref. 235027) was purchased from US Biological Life Sciences (Salem, USA). Pharma-grade sodium hyaluronate (M_w_ ∼ 620–1200 kDa; Ref. 4266107) and sterile ultrapure alginate Pronova SLG100 (M_w_ ∼ 120–250 kDa; Ref. 4202101) were purchased from Novamatrix (Sandvika, Norway). Acetone was purchased from Thommen-Furler AG (Rüti b. Büren, Switzerland). Poly(ethylene glycol) methyl ether (PEG; M_n_ ∼ 5 kDa, Ref. 81323), 3,6-dimethyl-1,4-dioxane-2,5-dione (lactide; Ref. 303143), tin(II)-2-ethylhexanoate (SnOct2; Ref. S3252), diethyl ether (Ref. 32203), calcium carbonate (CaCO_3_; Ref. 21069), δ-gluconic acid lactone (GDL, Ref. G4750) and ethyl acetate (Ref. 34858) were purchased from Sigma-Aldrich (Buchs, Switzerland). PBS was purchased from Thermo Fisher Scientific and endotoxin-free water was used for in vivo experiments (Thermo Fisher Scientific). Trazimera (Pfizer, USA) 420 mg and Herceptin SC (Roche) was provided by the pharmacy of the Institut de Cancérologie Strasbourg Europe.

### Methods

#### PEG-b-PLA synthesis

All flasks used were flame-dried and the glass condenser was dried at 105 °C. PEG (3.5 g; 0.7 mmol; 1 eq.) and lactide (11.2 g; 77.5 mol; 111 eq.) were dried overnight under vacuum by heating at 50 °C and gentle stirring. 30 mL of toluene were dried with freeze-pump-thaw cycles under argon atmosphere and added to the flask containing the reactants. Upon dissolution of lactide and PEG at 100 °C, SnOct_2_ (317 μL; 1 mmol; 1.4 eq.) was added to the reaction mixture, the temperature was raised to 140 °C, and the reaction was left under reflux for ∼6 h. The block copolymer was precipitated in ice-cold diethyl ether, filtered, and redissolved in DCM a total of three times. The resulting white solid was dried under vacuum. ^1^H-NMR spectra were acquired on a Bruker Avance III 400 (Bruker BioSpin GmbH) and the chemical shifts were reported relative to the solvent peak of CDCl_3_: δ = 7.26 ppm. ^1^H-NMR (400 MHz, CDCL3): δ = 5.29–5.05 (m, 237H), 4.39–4.21 (m, 3H), 3.76–3.52 (m, 555H), 3.37 (s, 3H), 1.64–1.41 (m, 725H). Molecular weight by ^1^H-NMR: PEG ∼6 kDa, PLA ∼18 kDa. GPC (poly(styrene)-*co*-divinylbenzene calibration): *M*_n_ = 34 kDa, *M*_w_ = 41 kDa, *Ð* = 1.23.

#### Nanoparticle synthesis

NPs were formed via batch nanoprecipitation whereby 140 mg of PEG-*b*-PLA were dissolved in 2.5 mL of acetone and added dropwise to 10 mL of dH_2_O under stirring. The residual acetone was evaporated under ambient conditions overnight. The NPs were purified and concentrated via ultrafiltration using Amicon Ultra centrifugation filters (MWCO: 30 kDa, 4500 rcf, 30 min) and washed with 9 mL of sterile PBS. This step was repeated a second time and the NPs were concentrated again via ultrafiltration at 4500 rcf for 50 min and resuspended to a final concentration of 20 wt%.

#### CD–PNP hydrogel formation

The whole study was performed with two hydrogel formulations: HA CD–PNP (2 wt% HA, 5 wt% NPs, 10 wt% αCD) and Alg CD–PNP (2 wt% alginate, 5 wt% NPs, 10 wt% αCD). For convenience, we describe the formation of 1000 mg CD–PNP hydrogels but this volume can be scaled up or down depending on the need. For HA CD–PNP hydrogels, 100 mg of αCD and 20 mg of HA were dissolved in 394 μL PBS (pH 7.4) within a plastic syringe. The syringe was sealed and allowed to equilibrate overnight. 250 mg of 20 wt% NPs and 236 μL PBS were prepared in a separate plastic syringe. The NP and polymer syringes were connected with a female-to-female luer-lock adapter (Cellink, Sweden), gently mixed for 5 min, allowed to equilibrate overnight at room temperature, and stored at 4 °C until further use. For trastuzumab-loaded formulation, the antibody was added instead of PBS to the NP-containing syringe. Alg CD-PNP hydrogels were prepared analogously as explained above.

For post-stabilization of the scaffold 2.8 mg of solid CaCO_3_ and 10 mg of solid GDL were added to a syringe. About half of the PBS that would have been used to dissolve alginate (394 μL) was implemented to form the calcium carbonate slurry. This means that if 200 μL PBS were added to the syringe containing alginate, then the remaining 194 μL PBS were used in a separate syringe to dissolve CaCO_3_ and GDL. To make the calcium-crosslinked hydrogels, supramolecular Alg CD–PNP hydrogels were formed and let equilibrate overnight and, right before injection, PBS was added to the syringe containing solid CaCO_3_ and GDL. The Alg CD–PNP and the calcium carbonate slurry syringes were connected with a female-to-female luer-lock adapter, mixed for 20 s, and used.^18^ Trastuzumab was loaded in the hydrogels using the FDA-approved pharmaceutical equivalent Trazimera by entrapping the antibody into the hydrogel network during formation. Trazimera was dissolved overnight in the NP suspension and subsequently mixed with the remaining components as described above. To prepare quantities lower than 300 mg, the same sequential mixing process was performed in Eppendorf tubes and, to ensure a correct ratio of components, the Ca^2+^ slurry was sometimes prepared on larger scales, well mixed, and only a fraction was pipetted into the syringe.

#### Rheology

The rheological characterization of the hydrogels was performed on a shear rheometer (MCR 502; Anton Paar; Zofingen, Switzerland) equipped with a Peltier stage and a covering hood. Hydrogels were tested at 25 °C using a parallel plate geometry with a diameter of 20 mm and a gap size of 0.5 mm. After loading the sample, an oscillatory interval (ω = 0.1 rad s^-1^, γ = 0.01 %, t = 30 min) was performed to reduce the loading history of the sample. The linear viscoelastic region was determined with dynamic oscillatory strain amplitude sweep measurements with constant angular frequency (ω = 10 rad s^-1^). The hydrogel response to different frequencies was probed with dynamic oscillator frequency sweep measurements conducted with constant strain amplitude (γ = 0.01 %) and a decreasing angular frequency (from 100 to 0.1 rad s^-1^). The ability of the CD–PNP formulations to flow and recover was characterized with dynamic oscillatory time sweep measurements with alternating low (γ_low_ = 0.1 %) and high (γ_high_ = 10 or 200 %) strain intervals at constant frequency (ω = 10 rad s^-1^). The shear-thinning properties of CD–PNP hydrogels were tested with rotational shear rate measurements. The shear rate, δγ/δt was increased from 0.01 to 100 s^-1^. The data was fitted to the following power-law model

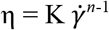

where *n* ∼ 1 for Newtonian, *n* > 1 for shear-thickening, and *n* < 1 for shear-thinning fluids.

#### Turbidity measurements

Polymer–trastuzumab complexation was confirmed via turbidity measurements. Solutions containing 2 wt% of polymer (HA or alginate) with or without trastuzumab (3 wt%) were formed in a 96 well plate. The solutions were left to equilibrate for 2 days on a shaking plate. Afterwards, the absorbance of the mixture as well as of pure PBS and PBS with 3 wt% trastuzumab was measured at 450 nm.

#### In vitro release of Trastuzumab

To study the in vitro release properties of trastuzumab from CD–PNP formulations, ∼100 mg of hydrogel containing 12 wt% trastuzumab were placed at the bottom of a 2.0 mL microcentrifuge tube. The experiment was started by gently adding 0.75 mL of 1X PBS and incubating the samples at 37 °C. At each timepoint, the supernatant was gently mixed once by aspirating and releasing 300 μL of supernatant. Subsequently, 300 μL were collected and replaced with fresh PBS. Quantification of released mAbs was accomplished through bicinchoninic acid assay (Pierce™ BCA Protein Assay Kits, Thermo Fisher Scientific) following manufacturer instructions. The absorbance of the supernatant was measured at 562 nm using a Hidex Sense and compared with calibration curves.

#### Drug release modeling

We used the Ritger-Peppas equation to analyze the early release kinetics to distinguish between an erosion- or a diffusion-based release.^21,22^ This equation describes both Fickian and non-Fickian release from the hydrogel, and is expressed as follows: M_*t*_/M_∞_ = *k* ⋅*t*^n^, Where, M_*t*_ and M_∞_ represent the released mass at time *t* and infinite time, respectively. *k* is a constant, and *n* is the power governing the release activity. In brief, a value of *n* between 0.42 and 0.5 indicates a purely diffusive system, while *n* > 0.5 suggests an anomalous release, often associated with gel degradation.

#### In vitro binding assays

The trastuzumab integrity post release was assessed by flow cytometry toward HER2+ breast cancer cell line HCC-1954. Firstly, trastuzumab was labeled with Cy5.5 prior to loading in the hydrogel formulation. After the in vitro release, supernatant was collected to assess the specificity of the trastuzumab. 1–2 × 10^5^ cells per condition were incubated with 100 μL of fluorescently-labeled trastuzumab in FACS buffer (PBS, FBS 2%, EDTA 2 mM) for 5 min at room temperature. The cells were washed three times with PBS amd centrifuged at 300 rcf for 5 min for each wash. MACSQuant® Analyzer 10 Flow Cytometer (Miltenyi) allowed data acquisition, and subsequent analysis was performed using FlowJoTM software (V. 10.8.2).

#### Dosage of trastuzumab in mouse serum

The quantification of plasma/serum levels of trastuzumab was conducted utilizing Time resolved-fluorescence linked immunosorbent assay (TR-FLISA) kits designed for mice or nonhuman primates (NHP), procured from Poly-Dtech (Strasbourg, France). Experimental procedures were conducted in accordance with the manufacturer’s guidelines. Briefly, microtiter plates were subjected to three washes with washing buffer, followed by the addition of 100 µL of standards and samples in duplicate. A standard curve for each mAb of interest was employed, utilizing dilution buffer as the zero standard (0 ng mL^-1^) and plasma collected from mice or NHP prior to administration as a negative control. Various plasma dilutions were examined for all mouse/monkey samples to establish the experimental setup. Subsequently, the plates were incubated with gentle shaking for 1 hour at 37 °C, followed by three washes. Next, 100 µL of the detection antibody, diluted in assay buffer, was added to each well and incubated with shaking for 1 hour at 37 °C. A final wash comprised six cycles (5 minutes incubation with shaking per wash) to minimize background interference. Following this, 200 µL of washing buffer was added to each well, and fluorescence readings were recorded at 535 nm (excitation at 340 nm) utilizing the time-resolved fluorescence (TRF) read mode of a SpectraMax ID5 (Molecular Devices), adhering to the TRF plate reader settings as outlined in the manufacturer’s instructions. Raw data were processed using the corresponding SoftMax Pro software, employing a 5-parameter logistic curve for analysis.

#### Pharmacokinetic parameters quantification

To extract the parameters for the pharmacokinetic profile, we employed an extravascular route of drug administration, characterized by a differential equation describing the drug concentration in blood over time, incorporating absorption and elimination rates: d*X*/d*t* = *K*_a_⋅*X*_a_ - *K⋅X*. Here, d*X*/d*t* represents the rate of change of drug concentration in blood, *X* signifies the amount of drug in the blood at time *t, X*_a_ denotes the amount of absorbable drug at the absorption site at time *t*, and *K*_a_ and *K* represent the first-order absorption and elimination rate constants, respectively. The pharmacokinetic parameters were derived from the solution of this equation. The half-life (*t*_1/2_) was determined using the formula *t*_1/2_ = ln(2)/K_a_. C_Max_, representing the maximum drug concentration, was obtained from the adjusted curve. The area under the curve (AUC) was calculated using the AreaXY function, integrating the curve with respect to elapsed time and the amount of drug in the blood. The entire analysis was conducted utilizing IGOR Pro (V.9.0, Wavemetrix).

## Author contributions

A.D. and M.W.T. conceived the project. G.B., S.B., E.A.G., and M.W.T. designed the hydrogel formulation with support from L.G.P. and L.L. on the ionic cross-linking design. G.B. carried out the hydrogel rheological characterization and the in vitro trastuzumab delivery experiments. V.M. supported hydrogel formulation selection and preliminary in vitro antibody release experiments. V.M. labeled trastuzumab with fluorophores and performed in vitro antibody binding experiments. G.B., A.D., C.M., and M.W.T. planned the in vivo pharmacokinetic experiments. G.B. and E.A.G. prepared the hydrogels for the in vivo experiments. G.B., C.M., S.H, G.J., and V.M. carried out the in vivo pharmacokinetic experiments, the histological analysis, and quantified the blood plasma concentrations of trastuzumab. O.L. and J.G. carried out the in vivo fluorescence analysis. L.C. and X.P. provided support and guidance on the experimental design. G.B., M.W.T., and A.D. wrote the manuscript with input from all authors.

## Acknowledgments

This work was supported by the Swiss National Science Foundation (200021_184697). We thank Chloé Guilbaud-Chereau for contributing to the initial discussions. Figures were drawn using elements of the Adobe Stock library.

