## Supplementary Figures for "Reinforced polymer–nanoparticle hydrogels for subcutaneous and sustained delivery of trastuzumab"

**Supplementary information**

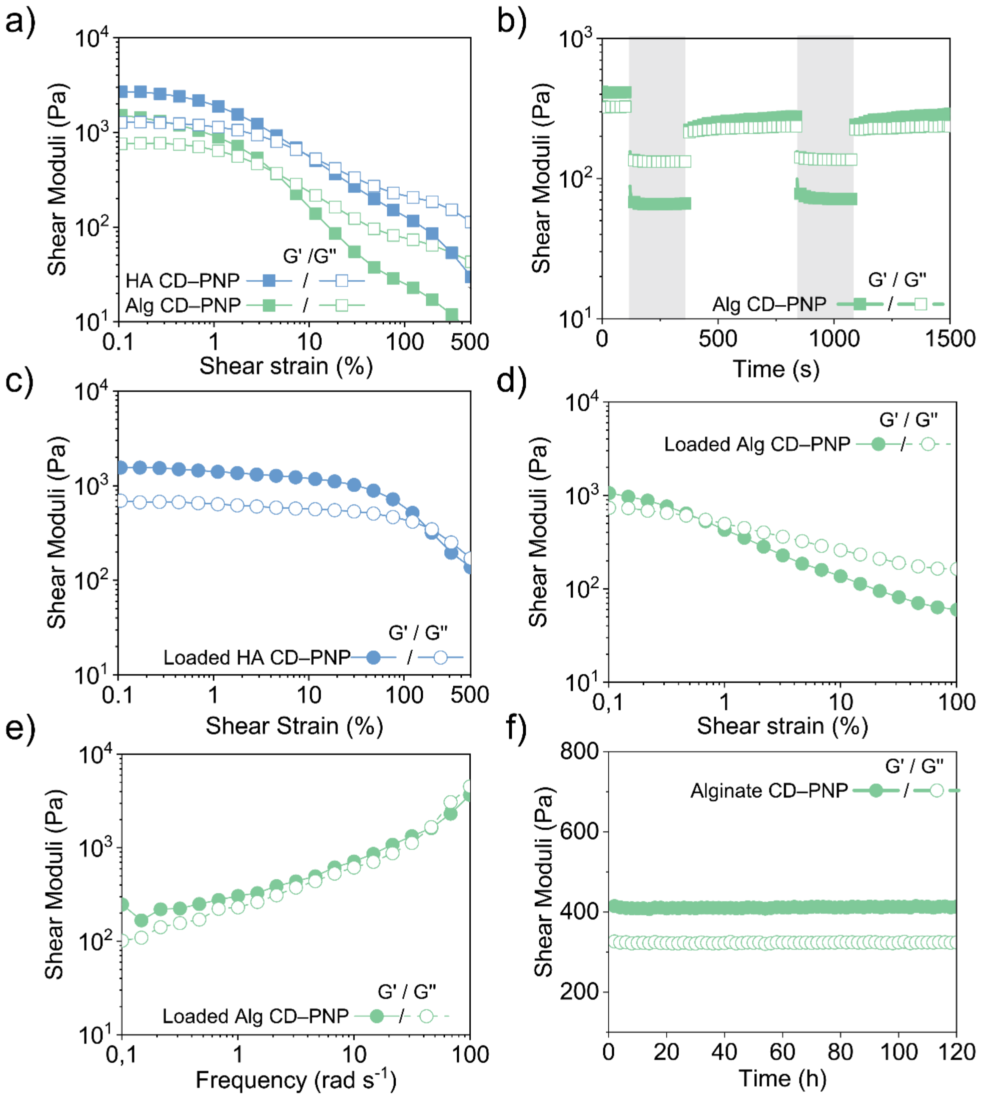


**Figure S1 HA and alginate CD–PNP hydrogel rheology.** a) Oscillatory strain sweep ($\omega$ = 10 rad s$^{-1}$) of HA CD–PNP (2 wt% HA, 5 wt% NPs, 10 wt% αCD) and Alg CD–PNP (2 wt% alginate, 5 wt% NPs, 10 wt% αCD). b) Alg CD–PNP hydrogels oscillatory strain recovery experiments comprised periods of high oscillatory shear strains ($\gamma$ = 200 %; $\omega$ = 10 rad s$^{-1}$) alternated to periods of low oscillatory shear strains ($\gamma$ = 0.1 %; $\omega$ = 10 rad s$^{-1}$). c) Oscillatory strain sweep ($\omega$ = 10 rad s$^{-1}$) of HA CD–PNP hydrogels loaded with 120 mg mL$^{-1}$ trastuzumab. d) Oscillatory strain sweep ($\omega$ = 10 rad s$^{-1}$) of alginate CD–PNP hydrogels loaded with trastuzumab. e) Oscillatory frequency sweep ($\gamma$ = 0.1 %) of alginate CD–PNP hydrogels loaded with trastuzumab. f) Time sweep ($\gamma$ = 0.1 %, $\omega$ = 10 rad s$^{-1}$ of non-loaded alginate CD–PNP).

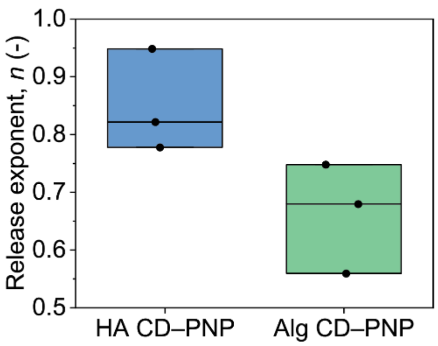


**Figure S2 Results from fitting to the Ritger-Peppas model.**[21,22] The first fraction of release was fitted to the Ritger-Peppas model. The determined release exponent, $n$, is 0.5 for Fickian diffusion, $0.5 < n < 1.0$ for non-Fickian transport, or $n \sim 1$ for zero-order release.
